# Deletion of *Cacna1c* (Ca_V_1.2) in D1-expressing cells elicits divergent sex-specific effects on aversive and spatial memories

**DOI:** 10.1101/2025.10.31.684545

**Authors:** Josiah D. Walsh, Diego Scala-Chavez, Andrew S. Lee, Arlene Martínez-Rivera, Anjali M. Rajadhyaksha

**Author notes:** These authors contributed equally (Josiah D. Walsh, Diego Scala-Chavez).

## Abstract

Dopamine signaling is critical for cognitive and emotional regulation and is implicated in multiple neuropsychiatric disorders. One downstream effector of dopamine is the L-type calcium channel Ca_V_1.2, encoded by the risk gene *CACNA1C*. Genome-wide association studies have consistently linked *CACNA1C* single nucleotide polymorphisms to schizophrenia, bipolar disorder, and related conditions. We previously showed that homozygous deletion of *Cacna1c* in dopamine receptor 1 (D1)-expressing cells enhances remote (30 days post-training) contextual fear memory in male mice. Here, we extend these findings by examining sex- and gene dosage-specific behavioral consequences of *Cacna1c* loss in D1 cells. We find that D1-*Cacna1c* deletion produces a sex- and gene dosage-dependent effect on fear memory. In males, homozygous loss of D1-*Cacna1c* heightens remote contextual fear at 30-days post-training, replicating prior findings, whereas partial loss had no effect. Cue-associated fear memory remained unaffected across genotypes. In contrast, females exhibited heightened contextual fear with both heterozygous and homozygous D1-*Cacna1c* loss at 24-hrs, 7-days, and 30-days post-training, indicating increased sensitivity to contextual aversive learning. Cue-associated fear memory was higher at 24-hrs but normalized at later time points in females. In the Water Y-maze, males with heterozygous or homozygous D1-*Cacna1c* loss showed impaired spatial memory at 7-days post-training, whereas females were unaffected. D1-*Cacna1c* deletion reduced locomotor activity selectively in females during the initial 5-mins of a 60-min session, with no genotype effects in males. Social interaction and anxiety-like behavior were unchanged across groups. Together, these findings highlight the interplay between dopamine receptor signaling and calcium channel function in shaping sex-dependent aspects of memory.

## 1 Introduction

Dopamine signaling is a central regulator of emotional, motivational, and cognitive processes, and dysregulation of dopaminergic circuits has been implicated in a broad range of neuropsychiatric disorders (Gholami & Mortezaee, 2025; Kim *et al*., 2015). Many therapeutics for psychiatric conditions target the dopamine D2 receptor; however, therapeutics for the dopamine D1 receptor, which plays a key role in cognitive processes (Goldman-Rakic *et al*., 2000; Kim *et al*., 2015) have not yet reached clinical use (Jones-Tabah *et al*., 2022). D1s are the most abundant dopamine receptor subtype in the human and rodent brain and are expressed in regions such as the striatum, cortex, hippocampus, and amygdala (Wei *et al*., 2018), where they regulate diverse aspects of cognition function. The downstream molecular mechanisms that mediate D1-dependent modulation of cognition remain incompletely understood. Defining these pathways is essential for clarifying how D1 signaling contributes to adaptive and maladaptive behavioral responses and inform new strategies for future therapeutic development.

A key molecular target downstream of D1 activation is the L-type voltage-gated calcium channel (LTCC), particularly the Ca_V_1.2 channel encoded by the *CACNA1C* gene (Hernández-López *et al*., 1997; Giordano *et al*., 2010). Converging evidence from genome-wide association studies (GWAS) has identified *CACNA1C* as a significant risk locus for multiple neuropsychiatric disorders, including schizophrenia, bipolar disorder, major depressive disorder, and autism spectrum disorders (Baker *et al*., 2023; Kabir *et al*., 2016, 2017). These conditions share disruptions in cognitive and affective regulation, overlapping with disturbance of D1-enriched brain regions where Ca_V_1.2 is expressed such as the striatum, prefrontal cortex, and hippocampus. In these regions Ca_V_1.2 channels govern activity-dependent neuronal plasticity and transcriptional regulation underlying learning and memory (Moosmang *et al*., 2005; Loganathan *et al*., 2024; Ireton *et al*., 2023; Tian *et al*., 2010). Work from our laboratory and others have shown that global and cell type-loss of *Cacna1c* disrupts cognition- and emotion-related behaviors (Kabir *et al*., 2017; Bavley *et al*., 2021; Lee *et al*., 2012; Loganathan *et al*., 2024). Notably, our previous work revealed that in male mice, selective homozygous deletion of *Cacna1c* in D1-expressing cells enhances contextual fear memory particularly at remote time points (Bavley *et al*., 2021).

Accumulating evidence suggests that sex is a critical biological variable influencing the neurobehavioral impacts of *Cacna1c* loss. In mice, Dao *et al*., (2010) reported sex-specific impairments in memory and social behavior following *Cacna1c* deletion in the forebrain and Klomp *et al*., (2022) reported sex-specific disruptions of motor performance, acoustic startle, and social behaviors. In rats, *Cacna1c* haploinsufficiency also produces sex-dependent alterations across social, affective, and cognitive behaviors (Bogdan *et al*., 2023; Braun *et al*., 2018; Kisko *et al*., 2020; Wöhr *et al*., 2021). Together, these findings position *Cacna1c* as a key modulator of behavioral domains disrupted in psychiatric disorders and suggest that its functional loss may produce distinct outcomes in males and females. These preclinical results parallel sex differences observed in *CACNA1C* risk allele carriers in human neuropsychiatric conditions (Strohmaier *et al*., 2013; Dao *et al*., 2010; Starnawska *et al*., 2016) and underscore the need to systematically evaluate sex- and gene dosage-dependent phenotypes in region- and cell-type-specific *Cacna1c* knockout models.

In the current study, we build upon our prior findings by testing whether the heightened remote contextual fear memory phenotype observed in male mice with homozygous loss of *Cacna1c* in D1-expressing cells also extends to females. We further examine whether heterozygous loss elicits behavioral effects, thereby revealing potential gene dosage sensitivity. In addition, we examine broader behavioral outcomes, including spatial learning and memory, locomotor activity, social interaction, and anxiety-like behavior, to define sex- and gene dosage-dependent behavioral outcomes of D1-specific *Cacna1c* loss.

## 2 Methods

### 2.1 Animals

Male and female conditional D1-*Cacna1c* knockout mice, in which *Cacna1c* was eliminated in D1-expressing cells, were generated by crossing homozygous *Cacna1c* floxed (*Cacna1c*^fl/fl^) mice with mice expressing Cre recombinase under the control of the Drd1 promoter (GENSAT; EY262 line) as previously described (Bavley *et al*., 2021). This resulted in D1-*Cacna1c* knockout (*Drd1a*^*Cre+*^;*Cacna1c*^fl/fl^, D1-*Cacna1c*^KO^) mice with homozygous deletion of floxed *Cacna1c* alleles, D1-*Cacna1c* heterozygous (*Drd1a*^*Cre+*^;*Cacna1c*^fl/+^, D1-*Cacna1c*^HET^) mice with heterozygous deletion of the floxed *Cacna1c* allele, and wild-type (*Drd1a*^*Cre+*^;*Cacna1c*^+/+^, D1-*Cacna1c*^WT^) littermates. All mice were >8 weeks old but <8 months old at the start of all behavioral experiments. Mice were maintained on a 12-hr light/dark cycle with *ad libitum* access to food and water. All procedures were conducted in accordance with the Weill Cornell Medicine Institutional Animal Care and Use Committee guidelines.

### 2.2 Contextual and Cued Fear Conditioning and Memory Recall

Fear conditioning was performed as previously described (Bavley *et al*., 2021) on day 1 and consisted of 2-min habituation followed by 5 trials (30-secs/trial) with increasing intertrial-intervals, where a tone (2.9 kHz, 38-dB) was played terminating in a shock (0.7 mA) within a rectangular chamber scented with (0.1%) peppermint odor and a rod floor. Context recall tests were performed on day 2, day 7 and day 30 in the original fear-conditioning chamber (Context A), whereas cue recall tests were performed on day 3, day 8, and day 31 in a distinct white cylindrical chamber (Context B) scented with lemon odor (0.1%) and a solid blue floor along with the presentation of the training tone. Behavior was recorded using a camera mounted above the soundproof box, and freezing was measured using FreezeView automated analysis (Coulbourn Instruments, Whitehall, PA).

### 2.3 Water Y-Maze

Mice were tested in the water Y-maze as previously described (Kabir *et al*., 2017). The Y-maze was filled with room temperature water, made opaque using non-toxic white paint. During training, mice performed 5 trials (60-secs/trial) with 60-sec inter-trial intervals. In each trial, mice were placed in an arm of the Y-maze and allowed to swim until they located a submerged platform (1-cm beneath water) in the goal arm. If a mouse was unable to locate the platform during a trial, it was removed from the water and placed on the goal platform for 15-secs before being removed from the Y-maze and placed in a heated cage, patted dry with paper towels, and allowed the 60-sec inter-trial interval rest before initiating the next trial. The latency to reach the platform was recorded manually using a stopwatch. The start arm and platform (goal arm) locations were randomized for each mouse but remained consistent across all training and testing sessions. Memory for the platform (goal arm) was assessed on day 2 and day 7 following training.

### 2.4 Locomotor Activity

Locomotor activity was evaluated over 60-mins using locomotor activity boxes (Med Associates, Fairfax. VT). Mice were placed in the center of a chamber (23.3cm x 23.3cm) at the beginning of the session. Activity was quantified as distance traveled in centimeters (cm) over 12 discrete 5-min time bins, which were collected with a computer-assisted locomotor activity monitoring software (Med Associates, Fairfax, VT).

### 2.5 Three-Chamber Social Approach

Mice were tested using the three-chamber social approach assay as described previously (Kabir *et al*., 2017). All testing was conducted in the three-chamber apparatus in a room with a ceiling-mounted camera for ANY-maze tracking. Two days prior to testing, age- and sex-matched C57BL/6J mice were individually placed under a wire cup in the left or right chambers and were observed for 10-min for disruptive behaviors such as bar-biting, circling, excessive grooming, or clinging to the side bars with all four paws. Only C57BL/6J mice that were deemed docile were used as novel mice during testing. On the test day, experimental mice were placed in the center chamber with the other chambers walled off for 5-min. Following this, the walls were removed, and the mice were provided an additional 5-min to explore the rest of the apparatus.

Next, mice were confined back to the center chamber while a novel mouse and novel object were placed in two barred cups on opposite chambers of the apparatus. The walls were then raised, and the mouse was provided 5-mins to freely interact with the stranger mouse and the novel object. Time spent in the contact zone (2-cm radius of the wire cups) was measured using AnyMaze automated analysis (Stoelting Co., Wood Dale, IL).

### 2.6 Elevated Plus Maze (EPM)

Mouse anxiety-like behavior was assessed using the EPM as previously published (Lee et al, 2012). Briefly, mice were placed in the center of the EPM and time spent in the open arms of the apparatus was measured. Behavior was recorded for 5 min using a camera mounted above the apparatus and time spent in each arm was obtained using the ANY-maze software (Stoelting Co., Wood Dale, IL).

### 2.7 Statistical Analysis

Statistical analyses were performed using R. For parametric analyses, 1-way ANOVA (II), 2-way ANOVA (III), or 2-way repeated measures (RM) ANOVA (III) were used. Planned contrasts for genotype comparisons on each test-day were performed with Bonferroni corrections for contextual and cue fear memory testing. For all other ANOVAs with significant effects, post-hoc tests were conducted with Bonferroni corrections. To test the assumption of a normal distribution, a Shapiro-Wilks test was conducted for all ANOVAs. To test the assumption of equality of variance, a Levene’s test (2-way ANOVAs) or a Mauchly’s test of sphericity (RM ANOVAs) was conducted. When parametric assumptions were violated, non-parametric analyses were performed. For 1-way ANOVA tests where the assumption of normality was violated, a Kruskal-Wallis H-test was performed, while a Welch’s *F*-test was used where the assumption of equality of variance assumptions was violated. For male and female water Y-maze training and memory testing datasets, Box-Cox or logarithmic transformations were performed to correct for normality violations prior to running 2-way RM-ANOVA. For 2-way RM-ANOVA tests where assumption of sphericity was violated, Greenhouse-Geisser or Huynh-Feldt corrections were applied for repeated measures effects. For significant effects from resulting from Welch’s *F*-tests, Games-Howell post-hoc corrections were applied. Sample sizes were based on previous work in the lab and are similar to other published work in the field. Sample sizes for individual experiments are reported in figure legends. Investigators were not blinded to genotype. Statistical significance was determined as a p-value < 0.05. All data displayed in figures are reported as mean ± standard error of the mean (SEM).

## 3 Results

### 3.1 Heterozygous D1-specific *Cacna1c* Loss is Sufficient to Heighten Contextual Fear Memory in Female but Not Male Mice

We previously reported that deletion of *Cacna1c* in D1-expressing cells in male mice heightens contextual but not cue fear responses at remote time points (Bavley *et al*., 2021). To replicate and extend these findings, we examined contextual and cue fear in male and female D1-*Cacna1c*^WT^, D1-*Cacna1c*^HET^, and D1-*Cacna1c*^KO^ mice (Figure 1). Mice underwent fear conditioning on day 1 consisting of five tone-shock pairings in context A (Figure 1A). Contextual fear memory was evaluated at long-term (day 2) and remote (day 7 and day 30) timepoints by returning mice to context A, whereas cue-associated fear was assessed in a distinct novel context B at long-term (day 3) and remote (day 8 and day 31) timepoints using the shock-paired tone.

**Figure 1.**
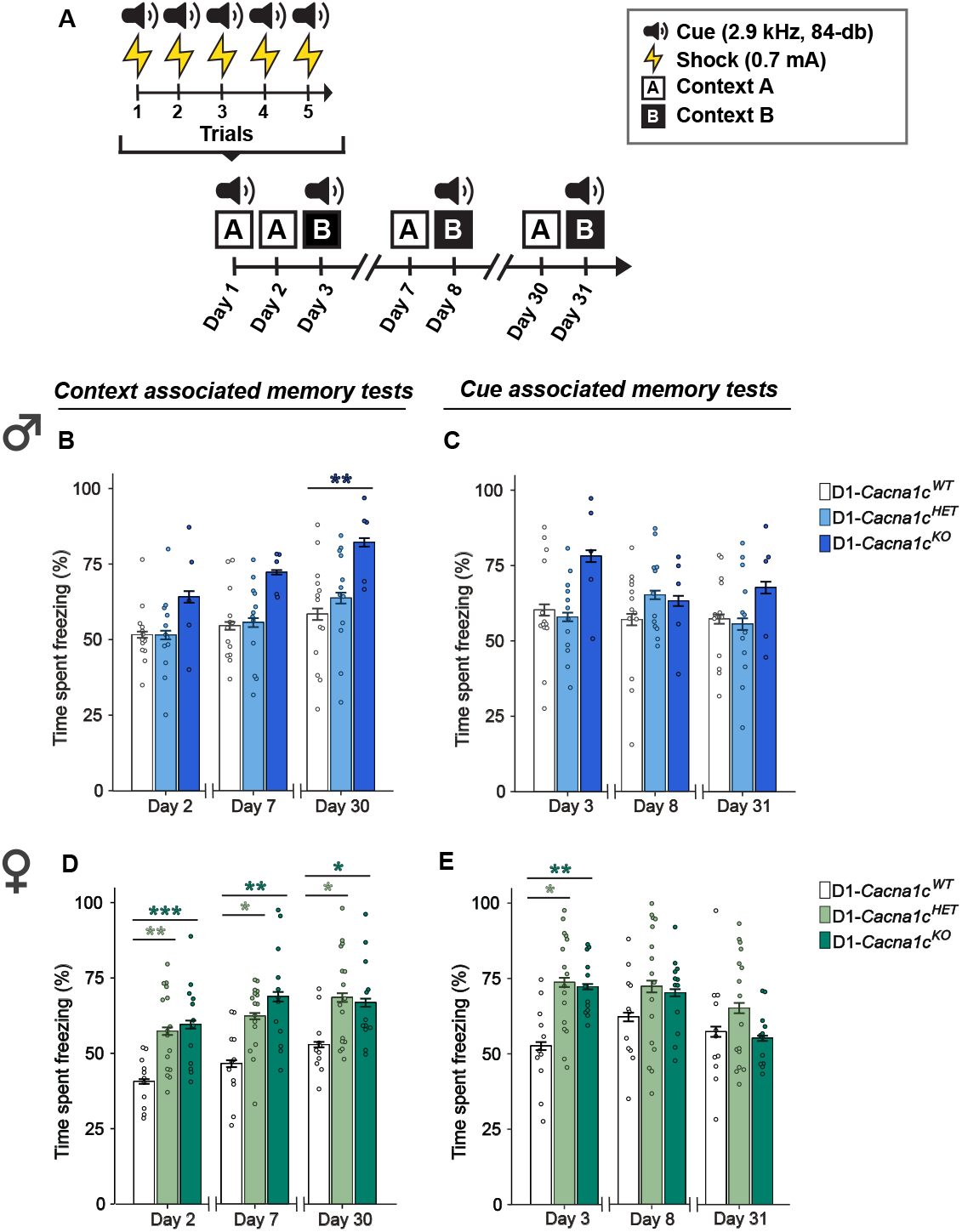
D1-specific *Cacna1c* deletion produces sex- and gene-dosage-dependent effects on fear memory. **(A)** Schematic of behavioral protocol to measure context- and cue-associated fear behavior. **(B)** D1-*Cacna1c*^KO^ males freeze significantly more than D1-*Cacna1c*^WT^ littermates during remote context memory testing on day 30 (\*\**p* = 0.0066, WT vs. KO, Bonferroni corrections). **(C)** D1-*Cacna1c*^HET^ and D1-*Cacna1c*^KO^ males display similar fear to cue across long-term (day 3) and remote (day 8 and day 31) memory testing as D1-*Cacna1c*^WT^ mice. **(D)** D1-*Cacna1c*^HET^ and D1-*Cacna1c*^KO^ females display significantly greater contextual fear than D1-*Cacna1c*^WT^ littermates during long-term (day 2) memory testing (\*\**p* = 0.0057, HET vs WT; \*\*\**p* = 0.0003, KO vs. WT, Bonferroni corrections) and remote memory testing on day 7 (\**p* = 0.016, HET vs. WT; \*\**p* = 0.0012, KO vs. WT) and day 30 (\**p* = 0.011, HET vs. WT; \**p* = 0.043, WT vs. KO). **(E)** D1-*Cacna1c*^HET^ and D1-*Cacna1c*^KO^ females display significantly greater cued fear than D1-*Cacna1c*^WT^ littermates during long-term memory testing (day 3) (\**p* = 0.011, WT vs. HET; \*\**p* = 0.003, WT vs. KO, Bonferroni corrections). Males: WT, *N* = 13; HET, *N* = 11; KO, *N* = 5. Females: WT, *N* = 11; HET, *N* = 16; KO, *N* = 12. Data are displayed as mean ± SEM.

In male mice, we replicated our previous findings, with D1-*Cacna1c*^KO^ mice showing significantly elevated contextual fear during remote memory testing (day 30) compared to D1-*Cacna1c*^HET^ littermates (Figure 1B: main effect of genotype: *F*_2, 27_ = 5.159, *p* = 0.013; main effect of day: *F*_2, 54_ = 7.577, *p* = 0.001, 2-way RM-ANOVA (III)). Conversely, there was no interaction or main effects of day or genotype during cue testing, suggesting similar cue fear memory in males (Figure 1C). In females, D1-*Cacna1c*^HET^ and D1-*Cacna1c*^KO^ mice exhibited significantly elevated contextual fear at both long-term (day 2) and remote (day 7 and day 30) memory timepoints (Figure 1D: main effect of genotype: *F*_2, 36_ = 13.038, *p* < 0.0001; main effect of day: *F*_2, 72_ = 7.561, *p* = 0.001, 2-way RM-ANOVA (III)). Additionally, during long-term (day 3) cue memory testing, D1-*Cacna1c*^HET^ and D1-*Cacna1c*^KO^ females showed greater fear memory than D1-*Cacna1c*^HET^ but genotype differences were absent during remote cue memory testing (day 8 and day 31) (Figure 1E: genotype × day interaction: *F*_4,72_ = 3.516, *p* = 0.011, 2-way RM-ANOVA (III); main effect of day: *F*_2,72_= 8.049, *p* = 0.0007). Together, these results reveal a sex- and gene dosage-dependent effects of *Cacna1c* in D1-expressing cells on conditioned fear memory, with females showing heightened sensitivity to fear memories, even with heterozygous *Cacna1c* loss.

### 3.2 Heterozygous and Homozygous D1-specific *Cacna1c* Loss Impairs Spatial Memory in Male but Not Female Mice

We previously reported that male mice with homozygous *Cacna1c* deletion in CaMKII-expressing cells exhibit learning and memory impairments in the water-based Y-maze task (Kabir *et al*., 2017). To determine whether similar learning and memory deficits occur following D1-specific *Cacna1c* deletion, we assessed male and female D1-*Cacna1c*^HET^ and D1-*Cacna1c*^KO^ mice alongside D1-*Cacna1c*^WT^ littermates.

Mice completed five 60-sec training trials to learn the location of a submerged goal platform (Figure 2). Memory performance was assessed as the average latency to reach the platform during retest on days 2 and 7. Both male (Figure 2B) and female (Figure 2E) D1-*Cacna1c*^HET^ and D1-*Cacna1c*^KO^ mice acquired the task at rates comparable to D1-*Cacna1c*^WT^ controls, with improvement from the first to the last trial across all males (Figure 2B: main effect of trial: *F*_4, 136_ = 4.568, *p* < 0.0001, 2-way RM-ANOVA (III)), although this trend was not observed at the genotype level, while females did not differ across trials (Figure 2E). Average latency across training trials was comparable across genotype in male (Figure 2C) and female (Figure 2F) mice.

**Figure 2.**
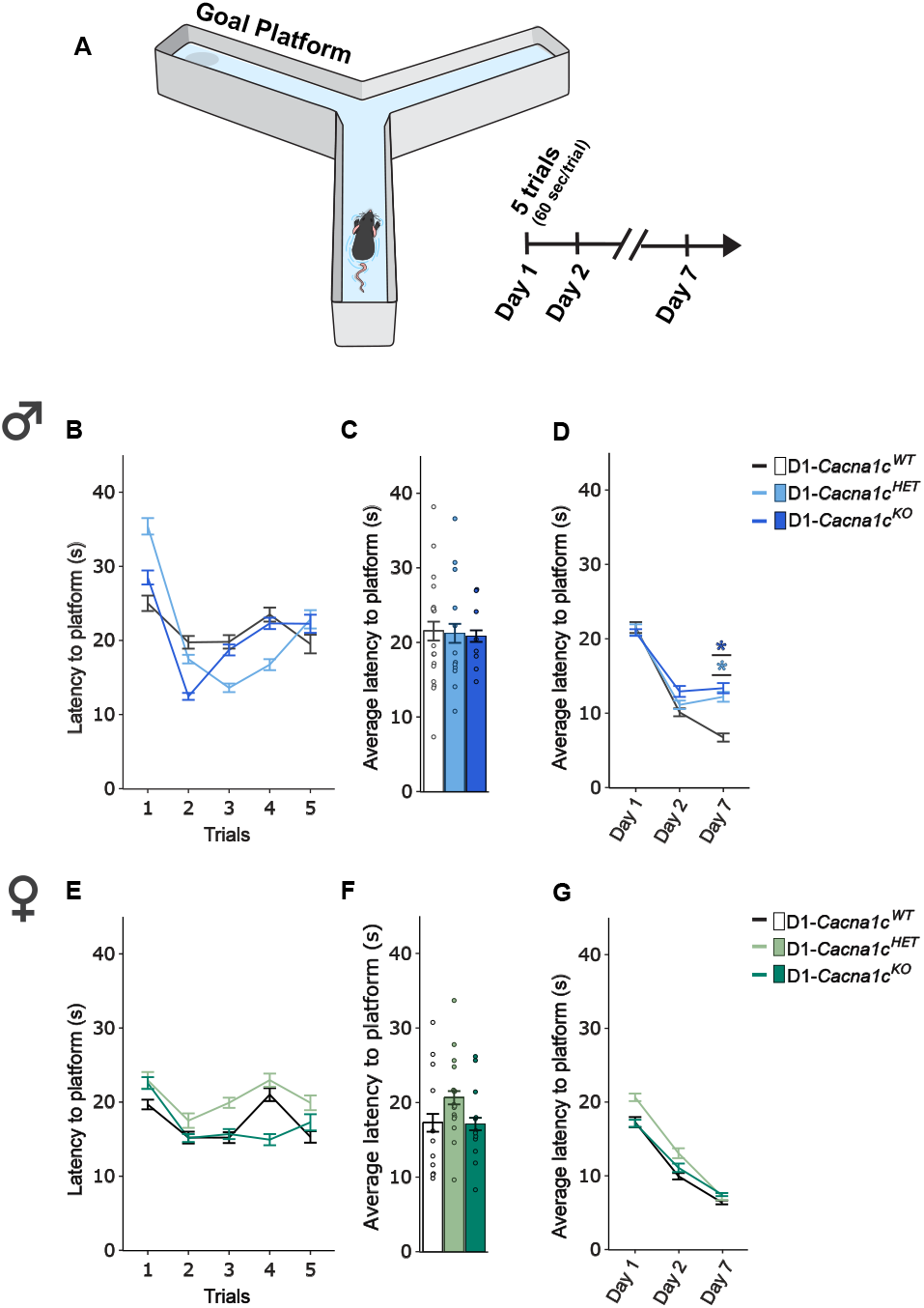
D1-specific *Cacna1c* deletion produces a sex-dependent effect on water Y-maze memory. **(A)** Schematic of behavior protocol to test learning, long-term and remote spatial memory. **(B)** During training, male D1-*Cacna1c*^HET^ and *Cacna1c*^KO^ display similar latency to reach submerged platform across trials as D1-*Cacna1c*^WT^. **(C)** Male D1-*Cacna1c*^HET^ and *Cacna1c*^KO^ did not differ in average latency to reach submerged platform during training compared to D1-*Cacna1c*^WT^. **(D)** Male D1-*Cacna1c*^KO^ and *Cacna1c*^HET^ display significantly higher latency to reach the submerged platform compared to D1-*Cacna1c*^WT^ littermates during remote memory (day 7) re-test (\**p* = 0.047, HET vs. WT; \**p* = 0.01, KO vs. WT, Bonferroni post-hoc corrections). **(E)** During training, female D1-*Cacna1c*^HET^ and *Cacna1c*^KO^ display similar latency to reach the submerged platform across trials as D1-*Cacna1c*^WT^. **(F)** Female D1-*Cacna1c*^HET^ and *Cacna1c*^KO^ do not differ in average latency to reach the submerged platform during training compared to D1-*Cacna1c*^WT^. **(G)** Female mice across genotypes display similar latency to reach the submerged platform across training, long-term (24-hr) and remote (7 day) memory testing. Males: WT, *N* = 13; HET, *N* = 11; KO, *N* = 8. Females: WT, *N* = 11; HET, *N* = 16; KO, *N* = 12. Data are displayed as mean ± SEM.

During memory recall, male D1-*Cacna1c*^HET^ and D1-*Cacna1c*^KO^ mice showed deficits at day 7 retest, requiring significantly more time to locate the platform compared to D1-*Cacna1c*^WT^ controls, while testing at day 2 was comparable across genotype (Figure 2D: genotype × day interaction: *F*_4, 68_ = 4.115, *p* = 0.005; main effect of day: *F*_2, 68_ = 44.749, *p* < 0.0001, 2-way RM-ANOVA (III)). In contrast, female mice showed comparable performance across genotypes on each test day (Figure 2G: main effect of day: *F*_2, 72_ = 71.835, *p* < 0.0001, 2-way RM-ANOVA(III). These results reveal sex- and gene dosage-dependent effect of D1-specific *Cacna1c* loss on spatial memory recall, with both partial and complete loss impairing performance in males while females are unaffected.

### 3.3 Homozygous D1-specific *Cacna1c* Loss Reduces Locomotor Activity in Female but Not Male Mice

To assess whether *Cacna1c* loss in D1-expressing cells affects basal locomotor activity, male and female D1-*Cacna1c*^WT^, D1-*Cacna1c*^HET^, and D1-*Cacna1c*^KO^ mice were placed in locomotor activity chambers and cumulative distance traveled was recorded for 60-mins. Locomotor activity was quantified as distance traveled across 5-min binned intervals.

In male mice, locomotor activity did not differ across geno-types throughout the session (Figure 3B), indicating that D1-specific *Cacna1c* loss does not affect basal activity. In contrast, D1-*Cacna1c*^KO^ females exhibited reduced locomotion compared to D1-*Cacna1c*^WT^ littermates (Figure 3D: genotype × time-bin interaction: *χ*^2^ = 0.05, *p* = 0.008, GG*ϵ*_13.88, 249.84_ = 0.631, *p* < 0.0001, 2-way RM-ANOVA (III)), indicating a sex-specific effect of *Cacna1c* loss on locomotor activity. Further analysis of the first 5-min of the session revealed that female D1-*Cacna1c*^KO^ mice traveled significantly less than D1-*Cacna1c*^WT^ and D1-*Cacna1c*^HET^ littermates (Figure 33E: main effect of genotype: *F*_2,23.2_ = 12.3, *p* = 0.0002, Welch’s *F*-test), revealing an early hypoactivity phenotype in D1-*Cacna1c*^KO^ females that was not observed in males (Figure 3C).

**Figure 3.**
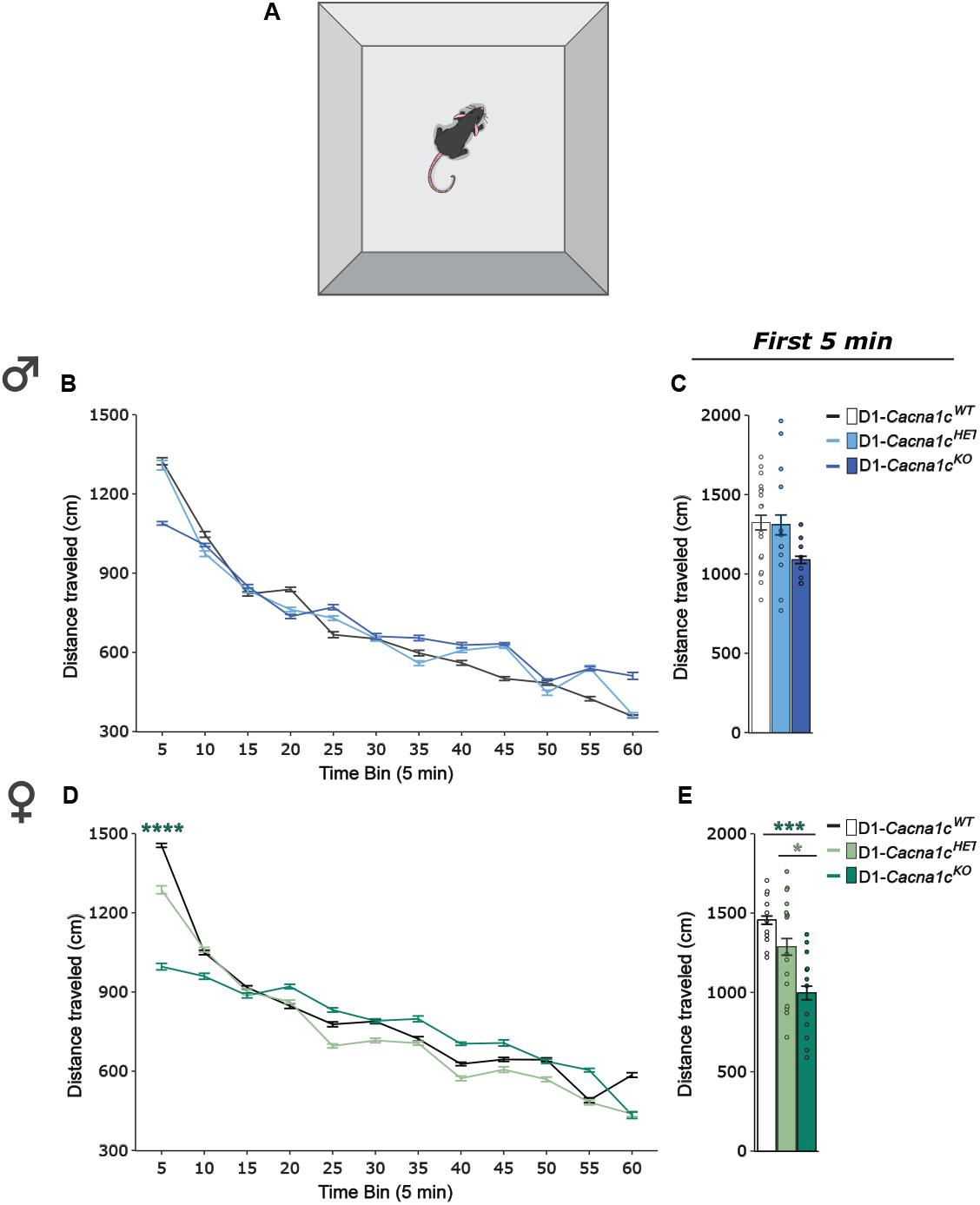
D1-specific *Cacna1c* deletion produces a sex-dependent effect on locmotor activity. **(A)** Schematic of locomotor apparatus. **(B)** Male D1-*Cacna1c*^KO^ and *Cacna1c*^KO^ display similar locomotor activity across time-bins as D1-*Cacna1c*^WT^. **(C)** Males across genotypes display similar locomotor activity in the first 5-min time-bin. **(D)** D1-*Cacna1c*^KO^ females travel a shorter distance during the first time-bin compared to D1-*Cacna1c*^WT^ littermates (\*\*\*\**p* < 0.0001, KO vs. WT, Bonferroni post-hoc corrections). **(E)** *Cacna1c*^KO^ females travel a significantly shorter distance than D1-*Cacna1c*^WT^ and D1-*Cacna1c*^KO^ littermates in the first 5-minute time-bin (\*\*\**p* = 0.0002, WT vs KO; \**p* = 0.041, HET vs KO, Games-Powell post-hoc corrections). Males: WT, n=13; HET, *N* = 11; KO, *N* = 8. Females: WT, *N* = 11; HET, *N* = 6; KO, *N* = 12. Data are displayed as mean ± SEM.

### 3.4 D1-specific *Cacna1c* Loss Does Not Alter Social Preference in Male or Female Mice

We previously reported that *Cacna1c* deletion in CaMKII-expressing cells impairs social preference in male mice (Kabir *et al*., 2017). To determine whether a similar deficit occurs with D1-specific deletion, we assessed social preference using the three-chamber social interaction test, where mice were first habituated to the apparatus for 10-mins and then were provided 5-mins to interact with a novel mouse and a novel object placed in barred cups in opposing chambers (Figure 4A).

**Figure 4.**
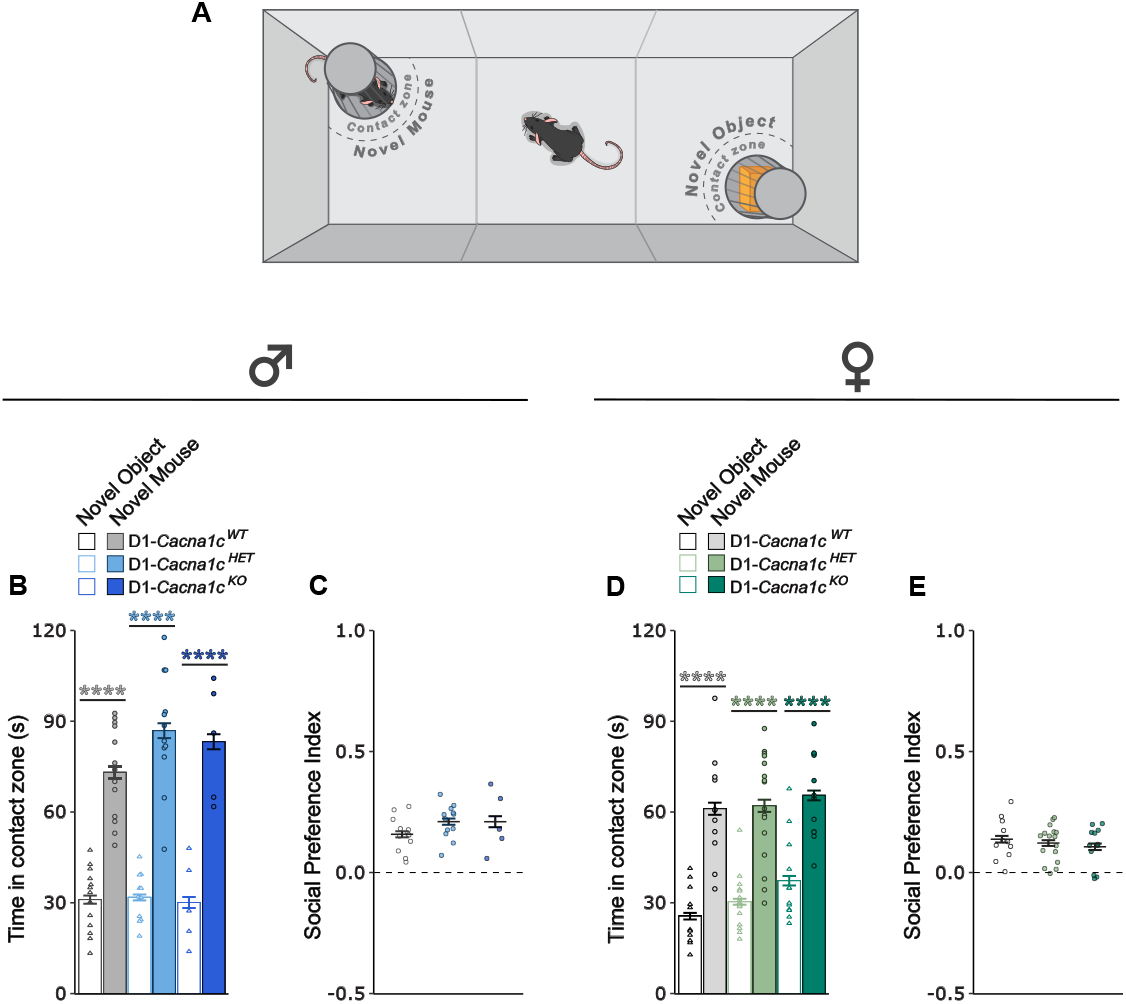
D1-specific *Cacna1c* deletion does not produce social preference deficits. **(A)** Schematic of apparatus used for the three-chamber social interaction behavioral test. **(B)** Male D1-*Cacna1c*^WT^, *Cacna1c*^HET^ and *Cacna1c*^KO^ mice spent significantly more time in the novel mouse contact zone than novel object contact zone (WT: \*\*\*\**p*< 0.0001; HET: \*\*\*\**p*< 0.0001; KO: \*\*\*\**p*< 0.0001, Bonferroni post-hoc corrections). **(C)** Males across genotypes did not differ across genotype for sociability index. **(D)** Female D1-*Cacna1c*^WT^, *Cacna1c*^HET^ and *Cacna1c*^KO^ mice spent significantly more time in the novel mouse contact zone than novel object contact zone (WT: \*\*\*\**p*< 0.0001; HET: \*\*\*\**p*< 0.0001; KO: \*\*\**p*= 0.0001, Bonferroni post-hoc corrections). **(E)** Females across genotypes did not differ across genotype for sociability index. Males: WT, *N* = 13; HET, *N* = 11; KO, *N* = 8. Females: WT, *N* = 11; HET, *N* = 16; KO, *N* = 12. Data are displayed as mean ± SEM.

All male mice displayed a significantly greater preference for the stranger mouse compared to the object (Figure 4B: main effect of chamber: *F*_1, 27_ = 136.260, *p* < 0.0001, 2-way RM-ANOVA (III)), with no genotype differences in social preference index (Figure 4C). Similarly, all female mice exhibited significantly greater preference for the stranger mouse compared to a novel object (Figure 4D; main effect of chamber: *F*_1,36_ = 85.527, *p* < 0.0001, 2-way RM-ANOVA (III)) and did not differ across genotypes on social preference index (Figure 4E). Thus, in contrast to CaMKII-specific deletion, D1-specific *Cacna1c* loss does not impair sociability in either sex.

### 3.5 D1-specific *Cacna1c* Loss Does Not Affect Anxiety-like Behavior in Male or Female Mice

We previously reported that *Cacna1c* deletion in CaMKII-expressing neurons induces anxiety-like behavior in male mice in the EPM (Lee *et al*., 2012; Kabir *et al*., 2017). To assess whether similar effects occur following D1-specific deletion, we measured anxiety-like behavior during a 5-min EPM test. Male and female D1-*Cacna1c*^WT^, D1-*Cacna1c*^HET^, and D1-*Cacna1c*^KO^ mice showed comparable exploration of the open arms (male, Figure 5B, C; female, Figure 5F, G) and open arm entries (male, Figure 5D; female, Figure 5H) with no genotype differences observed. Locomotor measures (total distance traveled) were unaffected by genotype in male mice (Figure 5E). However, in females, locomotor activity was significantly reduced in both D1-*Cacna1c*^HET^ and D1-*Cacna1c*^KO^ compared to D1-*Cacna1c*^WT^ littermates (Figure 5I: main effect of genotype: W_2, 23.612_ = 17.795, *p* < 0.0001, Welch’s *F*-test). Taken together, these findings indicate that *Cacna1c* deletion in D1-expressing cells does not alter anxiety-like behavior in either sex, though it may reduce locomotor activity in females.

**Figure 5.**
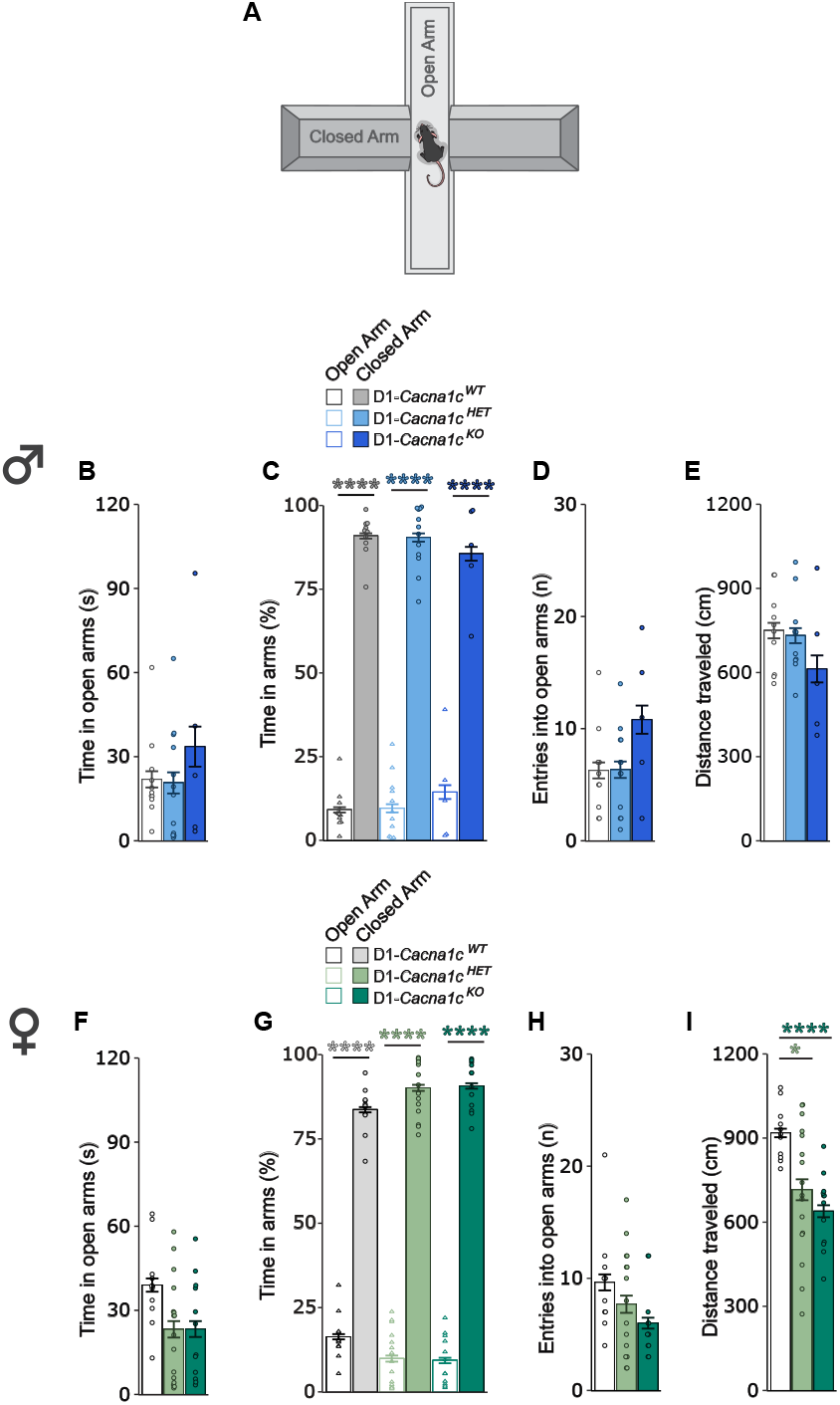
D1-specific *Cacna1c* deletion does not produce anxiety-like behavior. **(A)** Schematic of elevated plus maze used for anxiety-like behavior testing. **(B)** Male *Cacna1c*^HET^ and *Cacna1c*^KO^ mice spend similar time in the open arms to *Cacna1c*^WT^ mice. **(C)** Males regardless of genotype spent more time in the open arms compared to closed arms (WT: \*\*\*\**p*< 0.0001; HET: \*\*\*\**p*< 0.0001; KO: \*\*\*\**p*< 0.0001, Bonferroni corrections). **(D)** Males across genotypes perform comparable number of entries into the open arms across genotypes. **(E)** Males across genotypes traveled a comparable distance between genotypes. **(F)** Female *Cacna1c*^HET^ and *Cacna1c*^KO^ mice spend similar time in the open arms to *Cacna1c*^WT^ mice. **(G)** Females regardless of genotype spent more time in open arms compared to closed arms (WT: \*\*\*\**p*< 0.0001; HET: \*\*\*\**p*< 0.0001; KO: \*\*\*\**p*< 0.0001, Bonferroni corrections). **(H)** Females across genotypes perform comparable number of entries into the open arms across genotypes. **(I)** D1-*Cacna1c*^KO^ and D1-*Cacna1c*^HET^ females travel a significantly shorter distance than D1-*Cacna1c*^WT^ littermates (\*\*\*\**p*< 0.0001, WT vs. KO; \**p*= 0.014, WT vs. HET, Games-Howell post-hoc test). Males: WT, *N* = 11; HET, *N* = 12; KO, *N* = 5. Females: WT, *N* = 11; HET, *N* = 16, KO, *N* = 12. Data are displayed as mean ± SEM.

## 4 Discussion

In this study, we examined the behavioral consequences of selective *Cacna1c* deletion in D1-expressing cells, focusing on the influence of sex and gene dosage. Using fear conditioning, we found that *Cacna1c* loss produced distinct sex- and gene dosage–dependent effects on contextual fear memory. Females exhibited heightened fear responses to both context and cue at earlier time points, and to context alone at remote time points, even with heterozygous gene loss. In contrast, males required homozygous deletion to show enhanced remote contextual fear at day 30 post-training, without impacting early time points or on cue-associated fear. In the water-based Y-maze, heterozygous or homozygous D1-*Cacna1c* loss impaired remote spatial memory (day 7) selectively in males, while task acquisition and 24-hr memory was unimpaired, and performance across all days was comparable across genotype females. Indicating divergent roles of Ca_V_1.2 in aversive versus spatial learning across sexes. Locomotor activity was reduced in female mice with homozygous *Cacna1c* deletion, revealing an early hypoactivity phenotype, while males were unaffected. Neither heterozygous nor homozygous deletion of D1-*Cacna1c* impacted sociability or anxiety-like behavior in either sex. Together, these findings indicate that *Cacna1c* loss in D1-expressing cells exerts sex- and gene dosage-dependent behavioral outcomes, with females showing greater sensitivity to aversive memory and males displaying deficits in spatial memory.

Our finding that female mice with heterozygous loss of *Cacna1c* exhibit heightened sensitivity to remote aversive memories aligns with clinical evidence of higher PTSD prevalence and symptom persistence among women, such as re-experiencing and anxious arousal symptoms (Hanáková *et al*., 2024). Moreover, heritability estimates for mood- and stress-related disorders are higher in females (Amstadter *et al*., 2024). Consistent with this, female carriers of *CACNA1C* risk variants, such as rs1006737, display elevated risk for mood and affective disorders compared to male carriers (Strohmaier *et al*., 2013), and additional studies report sex-specific associations between *CACNA1C* SNPs and mood disorders (Dao *et al*., 2010). Beyond diagnostic risk, sex-dependent *CACNA1C* effects extend to functional recovery trajectories in schizophrenia-spectrum disorders (Strohmaier *et al*., 2013), stress-related transcriptional dysregulation (Dedic *et al*., 2018), and frontolimbic circuit activity (Zhu *et al*., 2019), typically with stronger effects in females. Epigenetic studies support this sexual dimorphism, with higher *CACNA1C* hypermethylation, predicted to lower *CACNA1C* expression, in females relative to males diagnosed with bipolar disorder (Starnawska *et al*., 2016). Collectively, these findings reinforce that *CACNA1C* influences neural and behavioral phenotypes in a sex-dependent manner, consistent with our preclinical finding of heightened sensitivity to aversive memories in females.

Our findings contrast with prior preclinical studies targeting other distinct cell populations and highlight a unique contribution of dopamine D1-expressing cells in contextual fear processing. For example, our previous work found that *CaMKII-Cre* driven *Cacna1c* deletion in excitatory neurons of male mice enhanced cue-associated fear memory (24-hrs post-training) without impacting contextual fear (Kabir *et al*., 2017; females not tested). This distinction underscores cell type specificity in Ca_V_1.2’s role in the type of aversive associative memories, with D1-expressing cells selectively mediating the persistence of contextual fear.

In contrast, water Y-maze testing revealed that remote spatial memory was disrupted by heterozygous or homozygous D1-*Cacna1c* loss only in males, aligning with evidence that cognitive impairments and memory retrieval problems are consistently observed in schizophrenia and bipolar populations (Guo *et al*., 2019; McCutcheon *et al*., 2023; Stefanopuolou *et al*., 2009; Torres *et al*., 2007; Bora *et al*., 2009; Gebreegziabhere *et al*., 2022), two disorders repeatedly associated with *CACNA1C* risk SNPs (He *et al*., 2014; Bigos *et al*., 2010; Nyegaard *et al*., 2010; Green *et al*., 2010; Ferreira *et al*., 2008). While the impact of *CACNA1C* variants on cognition and short-term recall of episodic and spatial memory has been repeatedly tested, its potential impact on long-term or remote spatial memory remains to be tested. For example, rs2007044 risk allele carriers exhibit more errors in spatial working memory (SWM) in a genotype dose-dependent manner (Cosgrove *et al*., 2017). Likewise, bipolar carriers of rs10466907, exhibited blunted improvements in cognitive performance following recovery from depression, despite comparable baseline performance (Lin *et al*., 2017). In contrast, rs1006737 carriers diagnosed with schizophrenia display impaired SWM, whereas bipolar carriers show minimal or opposite effects (Zhang *et al*., 2012), suggesting that distinct circuit-level vulnerabilities may underlie cognitive outcomes across disorders. Supporting this, Erk *et al*., (2010) found no episodic memory differences in bipolar rs1006737 carriers. Furthermore, male rs1006737 carriers exhibit reduced episodic memory and decreased activity in the anterior cingulate cortex and hippocampus (Erk *et al*., 2014). Together, these data support a modulatory role of *CACNA1C* in cognitive impairments and memory function, highlighting the importance of further exploring the intricacies across disorders and emphasizes the need for further clinical research on other forms of memory.

Our finding that heterozygous and homozygous loss of D1-*Cacna1c* impairs remote spatial memory (day 7 retest) in males, without impacting acquisition of the task or 24-hr memory performance, aligns with our prior work showing that homozygous D1-*Cacna1c* deletion in males disrupts remote spatial memory (day 30 retest) in the Morris water maze (Bavley *et al*., 2021). Although whether the effect extends to heterozygous deletion in males and testing in females remain to be assessed. Interestingly, females with homozygous loss of *Cacna1c* using the *Nex-Cre* line, which primarily targets post-mitotic pyramidal neurons in the cortex and hippocampus as well as mossy and granule cells in the dentate gyrus, produces a deficit in remote memory (day 7) not seen during training or 24-hr memory testing in the Morris water maze (Loganathan *et al*., 2024; males not tested). These results suggest overlap between cortical and hippocampal D1- and glutamatergic cell populations involved in remote spatial memory, while also indicating potential sex-dependent differences in their contribution. Notably, homozygous *Cacna1c* loss in CaMKII-expressing cells in male mice, which target a broader cell population, produces a learning deficit as well as memory deficits in the water Y-maze at both long term (24-hr) and remote (day 7) timepoints (Kabir *et al*., 2017). Females were not examined in this study. Similarly, a homozygous S1928 phosphorylation-deficient Ca_V_1.2 mouse model revealed impairments in learning and long term (24-hr) memory in males as well as females in the Morris water maze (Ireton *et al*., 2023). Together, these findings in mouse models indicate that *Cacna1c* loss disrupts remote spatial memory in a cell type- and sex-dependent manner, with deletions in broader cell types producing more widespread deficits in learning and long-term memory, highlighting the need to examine sex- and time-specific effects to fully understand Ca_V_1.2’s role in memory.

Mechanistically, our prior work showed that homozygous D1-*Cacna1c* deletion reduces hippocampal BDNF and adult hippocampal neurogenesis (AHN) in males (Bavley *et al*., 2021). A comparable mechanism may underlie female hypersensitivity, where heterozygous deletion reduces BDNF and AHN, which exacerbates aversive memory, manifesting at earlier time points than males. Supporting this, female but not male hemizygous carriers of the rs4765913 A-allele show reduced peripheral BDNF levels, and AA haplotype (rs476737–rs4765913) produces sex-opposite effects on BDNF expression, with females having lower and males having higher levels (Bastos *et al*., 2023). These findings suggest that *CACNA1C* variants may influence neurotrophic signaling and disproportionately enhance the aversive memories in females.

Sex differences in AHN arise from complex interactions among hormones, genetics, epigenetic regulation, and environmental stress, resulting in sex-biased cognition, stress reactivity, and psychiatric vulnerability (Galea *et al*., 2013; Gobinath *et al*., 2015; *Hodges et al*., 2022; Kempermann *et al*., 1997; Dao *et al*., 2010; Connolly *et al*., 2020). If reduced BDNF and AHN heighten sensitivity to aversive memories, this mechanism may preferentially affect contextual fear processing in females, while distinct hippocampal mechanisms mediate spatial deficits in males. Although the dorsal hippocampus contributes to both memory forms, females exhibit preferential engagement of the basolateral amygdala during fear retrieval (Keiser *et al*., 2017). Consistent with this, contextual fear activates CA1 neurons in males but suppresses them in females, who instead show increased basolateral amygdala activity (Colon & Poulos, 2022). The bed nucleus of the stria terminalis (BNST) also shows sex-dependent involvement, being required for contextual fear in males but not for cued fear in either sex (Urien & Bauer, 2022; Urien *et al*., 2021). Together, these findings suggest that sex-dependent regulation of aversive memory involves multiple interconnected circuits, with the amygdala and its afferents representing one potential critical node for mediating these differences.

Estrogen may further shape these effects by modulating Ca_V_1.2 function and downstream signaling. Estrogen directly enhances LTCC channel activity (Sarkar *et al*., 2008) and regulates L-type calcium channels through receptor-mediated mechanisms (Subbamanda & Bhargava, 2022). For example, 17β-estradiol increases AHN during spatial learning (McClure *et al*., 2013) and estrogens upregulate BDNF in hippocampal and dopaminergic neurons (Beyer & Karolczak, 2000; Ivanova *et al*., 2001; Zhou *et al*., 2005; Zhang *et al*., 2016; Barha *et al*., 2009; Barker & Galea, 2008). Thus, hormone-dependent regulation of Ca_V_1.2, BDNF, and AHN may contribute to the sex-specific memory phenotypes observed following D1-*Cacna1c* loss.

We also found that homozygous D1-*Cacna1c* loss reduces locomotor activity during the initial phase of exploration in females but not males, across open field and elevated plus maze testing. These results do not appear to be driven by an anxiogenic effect of D1-*Cacna1c* deficiency, suggesting a motoric rather than affective basis. Prior reports using *Syn1-Cre*-driven *Cacna1c* deletion noted female-specific motor deficits during rotarod testing; however, locomotor activity was unaffected (Klomp *et al*., 2022). Motor performance remains to be tested in mice with *Cacna1c* loss in D1 cells. This will be particularly important given the role of striatal dopamine and D1 receptors in motor learning (Giordino *et al*., 2018; Willuhn & Steiner, 2008; Phillips *et al*., 2024). In contrast to our findings, females with homozygous Nex-*Cacna1c* loss, exhibit hyperactivity in novel environments (Loganathan *et al*., 2024). Similarly, homozygous S1928A Ca_V_1.2 phosphorylation-deficient mice exhibit hyperactivity during 30-min locomotor activity testing, especially during later testing points, in male and female mice (Ireton *et al*., 2023). Additionally, while homozygous CaMKII-*Cacna1c* deficiency in males drives anxiety-like behaviors (Lee *et al*., 2012 & Kabir *et al*., 2017) and a similar phenotype is seen in male and female global heterozygous loss of *Cacna1c* (Lee *et al*., 2012), these effects were not shared in D1-*Cacna1c* males or females.

Finally, homozygous D1-*Cacna1c* loss does not alter sociability in male or female mice, distinguishing it from the social deficits observed with *CaMKII- or Syn1-Cre*-mediated deletions (Kabir *et al*., 2017; Klomp *et al*., 2022). In rats, global heterozygous loss of *Cacna1c* produces sex-specific social and affective changes. For example, juvenile males show reduced production of and approach toward 50-kHz ultrasonic vocalizations, a well-established index signaling positive affect, during rough-and-tumble play compared to wildtype rats, whereas heterozygous females display increased play and vocalizations, compared to wildtype littermates (Wöhr *et al*., 2021 & Kisko *et al*., 2021). This phenotype in juvenile females persists into adulthood, accompanied by increased grooming and reduced 50-kHz ultrasonic vocalizations (Redecker *et al*., 2019). However, in male but not female adult rats, *Cacna1c* haploinsufficiency reduces behavioral inhibition to danger signaling 22-kHz ultrasonic vocalizations played-back during exposure to predator urine (Wöhr *et al*., 2020). Together, these findings indicate that *Cacna1c* influences social behavior in a sex- and cell type-specific manner, likely independent of D1-expressing cells, though additional social behaviors remain to be tested, highlighting the importance of cell-type and sex in shaping its behavioral outcomes.

In summary, our findings demonstrate that *Cacna1c* loss in D1-expressing cells yields dissociable, sex-dependent behavioral outcomes; enhanced aversive memory in females and impaired spatial memory in males, while sparing social and anxiety-like behaviors. These results underscore the importance of Ca_V_1.2 in dopamine-modulated learning and highlight the need to dissect the circuit-level and hormonal mechanisms underlying sex-specific neuropsychiatric behavioral vulnerability linked to *Cacna1c*.

## Acknowledgments

This work was supported by grants from the NIH to A.M.R. (R01MH125006, R01DA053261, R01DA054368, R01MH118934). A.S.L. was supported by a Weill Cornell Medicine Clinical & Translational Science Center Pre-doctoral Training Award (TL1TR002386) from the National Center for Advancing Translational Sciences.

